# A hydrodynamic antenna: novel lateral line system in the tail of myliobatid stingrays

**DOI:** 10.1101/2024.10.08.616864

**Authors:** Júlia Chaumel, George V. Lauder

## Abstract

Eagle rays, cownose rays, and manta rays (family Myliobatidae) have a slender tail that can be longer than the animal’s body length, but its function and structure are unknown. Using histology, immunohistochemistry, and 3D imaging with micro-CT scans, we describe the anatomy and function of the tail in *Rhinoptera bonasus*, the cownose ray. The tail is an extension of the vertebral column with unique morphological specializations. Along the tail behind the barb, vertebral centra are absent and neural and hemal arches fuse and form a solid mineralized structure that we term the caudal synarcual, which imparts passive stiffness to the tail, reducing bending. Two lateral line canals connected to an extensive tubule network extend along both sides of the tail. Tubules branch from the lateral line canal toward the dorsal and ventral tail surfaces and open to the surrounding water via pores. A continuous neuromast is located within each lateral line canal, maintaining an interrupted structure along the entire tail. The complex lateral line mechanosensory system in the tail of *R. bonasus* supports the hypothesis that the tail functions like a hydrodynamic sensory antenna and may play an important role in their behavioral and functional ecology.

## 1. Introduction

Batoids (rays and skates) constitute the largest group of cartilaginous fishes, which also includes sharks and chimaeras [1]. Batoids are characterized by a dorsoventrally flattened body and are distributed worldwide, inhabiting a wide range of depths (up to 3000m), primarily in benthic and epibenthic ecosystems [1, 2]. The only batoids with a pelagic lifestyle are stingrays of the family Myliobatidae (eagle rays, cownose rays, and manta rays) comprising 19 species [1]. Their adaptation to a pelagic lifestyle is reflected in their distinctive body morphology, locomotion style, and how they interact with and sense their environment [3, 4]. Myliobatids present a triangular body shape with enlarged pectoral fins that oscillate vertically to move the animal forward, allowing them to swim effectively at different speeds and migrate long distances. [3, 5]. They also possess a well-developed lateral line system on the head and body, as well as an electrosensory system which is predominantly located on the ventral body surface in those species that detect prey buried in the sea-bottom [6–9]. However, as pelagic species, it remains unclear how myliobatids use these senses to detect stimuli, such as predators approaching from behind or localizing conspecifics during the formation of large schools, characteristic of many species in this group [10].

In addition to their specialized body shape, myliobatids are characterized by having a long and slender tail (often referred to as the whip-tail) which extends caudally from the body and supports a dorsal barb located at the tail base (Fig. 1A) [1]. The whip-like tail is an exclusive feature of myliobatiforms and is absent in other batoids and other chondrichthyans [1]. Unlike most bony fishes and sharks [11, 12], myliobatids do not use their tails as a propulsive structure. Although low amplitude movement of the myliobatid tail occurs during steady locomotion and maneuvering, the tail generally maintains a rostro-caudal orientation behind the body during swimming [3]. This positioning suggests the tail may have a sensory role, potentially detecting approaching predators or providing information on water flow and body position. However, no information on the anatomy and function of myliobatid’s tail is currently available to corroborate this proposed sensory function.

**Fig. 1.**
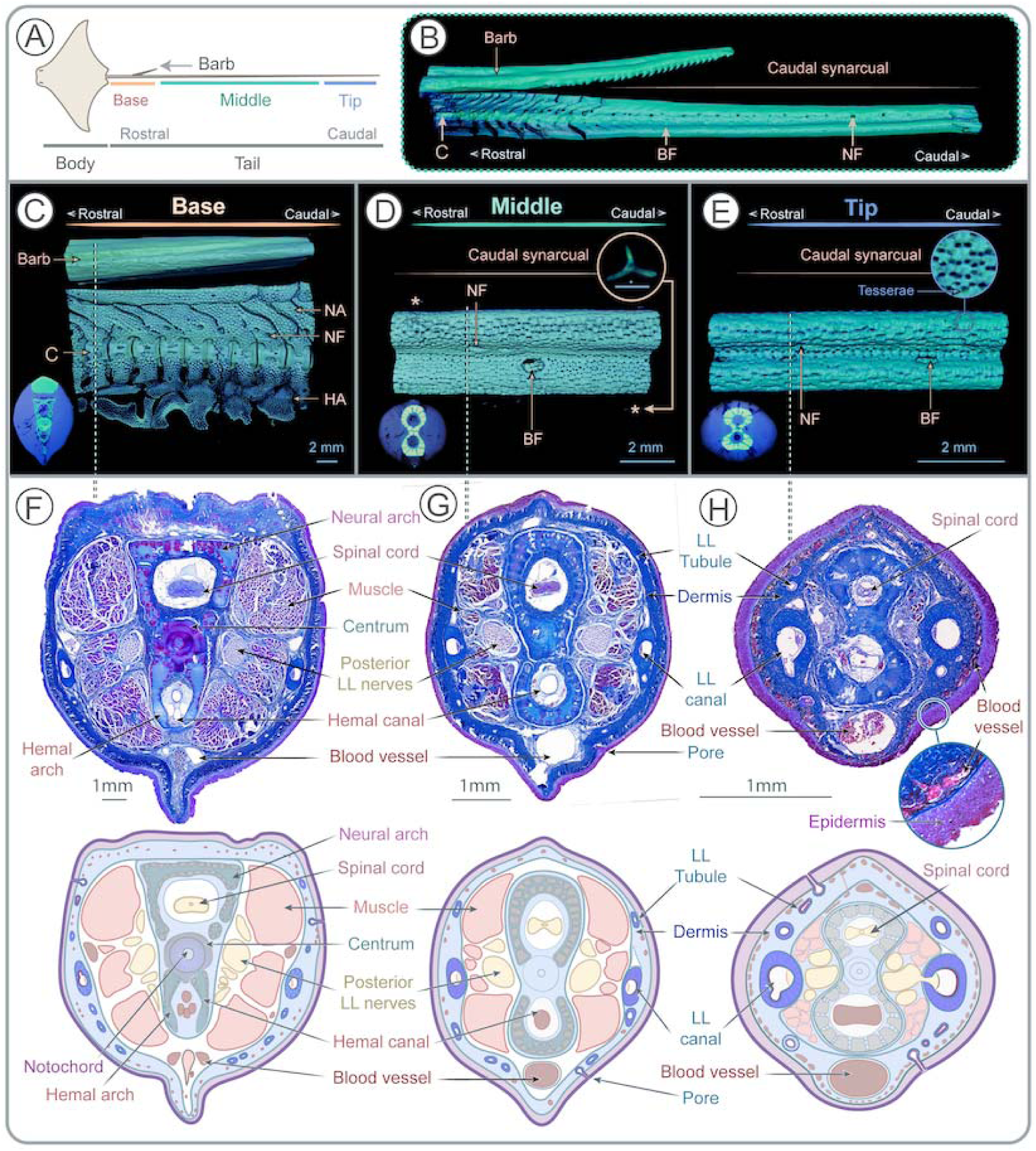
Tail anatomy. **A)** Tail division into base, middle and tip zones. **B)** µCT scan showing the transition from the segmented vertebrae to the caudal synarcual, with blood and nerve foramina following similar locations as in the anterior vertebrae. **C)** µCT scan of the tail base (more proximal to the body), characterized by a vertebral column with separated vertebrae units with neural arches, hemal arches, and by a well-defined centrum. **D)** µCT scan of the middle zone with caudal synarcual covered by tesserae and no centra. This is the only zone with denticles. **E)** µCT scan of the tip zone (more distal to the body) showing the continuity of the caudal synarcual until the end of the tail. **F–H)** Cross-sections of each tail zone stained with Mallory trichrome with an illustrative guide to the key structures below. Note the variation in vertebral morphology and the consistent presence of the spinal cord and the hemal canal. Muscles (red) are predominant at the base and are gradually reduced in the middle and tip zones. In all zones, posterior lateral line nerves (lilac color) run parallel to the vertebral column, the thick dermis that surrounds the tail (blue) and the epidermis (pink). The lateral line canals (both at each side of the tail) and the tubules are embedded in the dermis, while surface pores are located in the epidermis. **Abbreviations:** Blood foramina **(BF),** Centrum **(C)**, Denticles (**\***), Hemal arch **(HA),** Lateral line (LL), Neural arch **(NA)**, Nerve foramina **(NF)**.

The overall aim of this study is to clarify the sensory function of the tail in *Rhinoptera bonasus* (cownose rays), which may provide insights into a more comprehensive understanding of the biological function of the myliobatid tail. Our approach involves a combination of morphometric, gross anatomical, 3D imaging, and histological analysis using tails of cownose rays, a species with a well-documented ecology that occupies diverse ecological niches [13, 14]. Specifically, we describe morphological features of the tail and how they vary along its length, characterize the remarkable 3D structure of the sensory organs of the lateral line system that extends along the entire tail, and propose a potential function of the tail lateral line as a hydrodynamic filtering system. A detailed understanding of tail anatomy in cownose rays lays the groundwork for future comparative studies of tail anatomy in other ray species with diverse ecologies, as well as behavioral and biomechanical analyses of tail use and function during locomotion.

## 2. Methods

### (a) Sample collection

A total of 8 specimens of *Rhinoptera bonasus,* including males and females of different sizes, were used in this study (Supp. Mat. Table 1). Animal sizes were defined by disc width (DW), described as the distance between the tips of the pectoral fins. For gross morphometrics and comparative analysis, six individuals from the ichthyology collection at the Museum of Comparative Zoology of Harvard University were used (Supp. Mat. Table 1). These samples are stored at the collection in 70% ethanol. For fine-scale histological characterization, two fresh specimens were obtained from Ripley’s Aquarium of Myrtle Beach, South Carolina (DW=81cm, male), where tails were collected immediately after the natural death of the animal, and Havenworth Coastal Conservation during GULFSPAN survey in Florida (DW=50cm, male), where the animal was euthanatized following the ethical procedure under U.S. Fish and Wildlife Permit #SAL-1918-SRP (Supp. Mat. Table 1). Entire tails from fresh specimens were cut into small pieces (around 1cm length for histology and 2-4cm length for X-ray micro-computed tomography, µCT) immediately after animal’s death and fixed in paraformaldehyde 4% in phosphate buffered saline (PBS) during 6 hours at room temperature. Tails were divided into three zones: the base (more proximal to the body), a zone located between the end of the pelvic fins to the end of the barb, a middle zone located between the end of the barb and the tip region, and the tip zone (more distal to the body) corresponding to the last 15% of the tail (approximately 5cm in length for our samples) (Fig. 1A). As the division of the tail zone was defined based on anatomical features, each zone was characterized by different lengths and, therefore, number of samples analyzed. For histological and immunofluorescence analysis, 10 samples were analyzed per animal (3 from the base, 4 from the middle, and 3 from the tip), while µCT scans involved 5 samples per animal (2 from the base, 2 from the middle, and 1 from the tip zone).

### (b) X-ray micro computed tomography (µCT)

Samples were transferred into increasing ethanol concentrations (10%, 30%, and 50%) up to 70% ethanol. High-resolution µCT images were obtained using a Bruker SkyScan 1273 (70 kV and 114 uA). Image pixel size ranged between 4.9 and 13 µm. Samples were post-stained in 0.3% phosphotungstic acid (PTA, AMRESCO, Fountain Parkway Solon, OH) following [15] for over a month (changing PTA solution every two weeks) to assure a complete staining of all tissues. Once the staining was complete, PTA-stained samples were transferred to 70% ethanol and re-scanned. Images were reconstructed with Bruker Nrecon software and visualized, segmented, and analyzed using Amira-Avizo visualization software 2023.11 (ThermoFisher).

### (c) Histology and Immunofluorescence

After fixation, samples were decalcified using Epredia Decalfying Solution (ThermoFisher) for 48 hours and dehydrated in ethanol solutions of increasing concentrations, ending with immersion in 100% ethanol. Samples were transferred into 100% xylene, embedded in paraffin, cut in 5–7 µm thick sections with a microtome, mounted in VistaVision^TM^ HistoBond^®^ Premium Adhesion Slides (VWR International, LLC, Radnor, PA) and left to dry on a warming tray at 37°C overnight.

For histology, sections were stained with hematoxylin-eosin (HE) with Orange G and Mallory trichrome (protocols customized from [16]). Hematoxylin (purple) is a basic stain that binds to acid structures (e.g. nucleus), while eosin (pink) and Orange G (orange) are acidic stains with affinity for basic structures (e.g. cartilage matrix). The combination of both acidic stains helps to enhance different connective tissues by providing additional contrast. Mallory trichrome was used to identify muscle (red to orange), nervous system (lilac), collagen (dark blue), dense cellular tissue (pink to purple), mucus and connective tissue (blue), myelin and red blood cells (pink to red), and nuclei (red). Histology-stained samples were imaged using a Keyence digital microscope VHX-600 (Keyence Corporation, Japan) and a Leica light microscope (Leica Microsystems GmbH, Germany). Images were analyzed using Fiji 2.0.0.

For immunofluorescence staining, manufacturer protocol for the specific antibody was used. For antigen retrieval, samples were incubated in 10 mM sodium citrate buffer with 0.05% Tween 20 (pH 6) during 6 hours at 60°C. This method preserved the integrity of samples and the antibody signal. Samples were permeabilized in PBS 0.025% Triton X-100 and blocked with 5% bovine serum albumin in PBS (BSA; Sigma Aldrich) for 2 hours at room temperature. The primary antibody β-III-tubulin (1:500; Abcam, Cat# ab18207) was used as a neuronal marker and it was chosen due to its previously demonstrated reactivity with catshark tissues [17]. Samples were incubated with the primary antibody in 1% BSA in PBS overnight and then incubated during 1 hour at room temperature with a secondary antibody goat anti-mouse IgG H&L Alexa Fluor 488 (1:500; Abcam, Cat# ab150113) diluted in 1% BSA and then rinsed in PBS. Some tissue structures displayed an autofluorescence signal at 488nm. To avoid interference with the antibody signal, samples were stained with 0.1% Sudan Black B (Sigma Aldrich) following protocols established by [18, 19]. Samples were washed in PBS to remove the excess stain and mounted with SlowFade^TM^ Diamond antifade mountant with DAPI (ThermoFisher). This mountant with DAPI added additional autofluoresnce signal in the 405 channel, especially in the dermis. However, this did not affect the visualization of the structures of interest. Images of sections stained with immunofluorescence were acquired using a Zeiss LSM 980 confocal microscope (Carl Zeiss GmbH, Germany) and a Leica DM R fluorescence microscope (Leica Microsystems GmbH, Germany). Images were analyzed using Fiji 2.0.0.

Negative controls were performed (Supp. Mat. Fig. 2). The autofluorescence signal present when the sample was excited with the 405 wavelength laser was more pronounced by increasing image brightness. Such signal was used to obtain grey-scale images that showed the general tissue anatomy and were used as reference for the position in the section of the structures of interest following [20].

### (d) Measurements and analysis

Morphometrics of specimens’ body (total length, body length, and disc width) and tails (tail length and diameter) were measured *in situ* (Supp. Mat. Table 2). The lateral line system was visualized in PTA-stained samples with µCT and posteriorly segmented using Amira Avizo, obtaining a 3D model of the lateral line canal (LLC), tubules and pores (measured morphological parameters summarized in Supp. Mat. Table 2). The branching pattern and tubule divisions were characterized using Amira Avizo. Diameters and lengths of the LLC, tubules, and pore openings were measured using a combination of Amira-Avizo and Meshlab. The sum of the cross-sectional area of pores from the group of pores connected to the same primary tubules (termed *pore cluster* in Results section) was compared with the cross-sectional area of the LLC and primary tubules. Pore density was measured by counting the number of pores per cm of tail for each tail zone in all specimens (Supp. Mat. Table 1). Statistical analyses were performed to identify correlations between disc width and tail length, diameter variations of the LLC and pores, and variations in pore density across tail zones and size of the animals. Analyses were performed using linear models, ANOVA, and Tuckey poshoc tests using R 2023.12.0+369.

## 3. Results

### (a) Tail morphometrics

*Rhinoptera bonasus* is characterized by having a long and slender tail with a cone-like tapering geometry (i.e. tail diameter decreases from the base toward the tip). Larger animals have larger and wider tails; however, the tail does not scale proportionally with the animal’s size (measured as disc width) (Sup. Mat. Fig. 1A). In smaller animals, tails are 1.7x larger than the disc width (1:1.7 ratio), while in larger animals the tail has a similar length to the disc width (1:1 ratio) (Sup. Mat. Fig. 1B).

### (b) Tail anatomy

Micro CT data show that the tail contains an extension of the vertebral column which elongates from the relatively thick base toward the thinner tip of the tail (Fig. 1). However, the morphology of the vertebral column varies across different tail zones. At the base (from the tail attachment location to the posterior end of the barb; Fig. 1A–C), the vertebral column is similar in organization to the vertebral column of the body, composed of a series of repeated vertebral units characterized by a centrum (an areolar calcified hourglass-shaped structure) that surrounds the notochord; by dorsal neural arches that surround the spinal cord; and by ventral hemal arches that protect the caudal artery and vein (Fig. 1B,C). Both neural and hemal arches are covered by tesserae (mineralized cartilage) which is perforated by foramina that allow nerves and blood vessels to connect the caudal artery, veins, and spinal cord to the surrounding tissues. After the posterior end of the barb and proceeding distally to the middle and tip zones, the centra is absent and the neural and hemal arches fuse, forming a solid element that extends to the tip (Fig. 1D, E). This solid structure is also covered by tesserae and contains the spinal cord (dorsally) and artery and caudal vein (ventrally), both connected to the surrounding tissues through nerve and blood foramina. The spinal nerves connect the spinal cord to the posterior lateral line nerves (PLLN) that run parallel to the vertebral column in a caudal direction (Fig. 1F–H stained in lilac, Supp. Mat. Fig. 2). The vertebral column and PLLN are both surrounded by muscles; however, these muscular bundles are more predominant at the base of the tail and almost absent at the tip (Fig. 1F–H, stained in red).

Skeletal elements and muscles are further surrounded by a thick dermis (Fig. 1F–H). The dermis is further surrounded by an epidermis, formed of several layers of columnar cells (Fig. 1H). Embedded within the epidermis, and exclusively in the middle zone, the tail is covered by small (∼100µm) spiny denticles (Fig. 1D). Dorsally, the epidermis also contains melanocytes.

### (c) The lateral line system

The lateral line system is found embedded in the dermis and extends on each lateral side along the entire tail. Each lateral line system consists of a lateral line canal (LLC) connected to an extensive network of tubules (Fig. 2, Supp. Mat. Video 1; Supp. Mat. Fig. 3). The tubule network presents a tree-like topology, where primary tubules extend from the LLC in both dorsal and ventral directions (Fig. 2A–C, Supp. Mat. Fig. 3). Primary tubules are the initial and larges tubules that branch from the LLC. Primary tubules further divide into secondary tubules, which then divide again into tertiary tubules, which continue dividing until reaching a maximum of eight divisions (Fig. 2A–C). Tubules eventually open to the environment via pores (Fig. 2, Fig. 3). On the skin surface, pores are organized into groups of 15-20 pores, where each pore group derives from a single primary tubule (Fig. 2A–C, Fig. 3A). We refer to these groups of pores as “pore clusters” (Fig. 2A–C, Fig. 3A–B). With each division, the diameter of the tubules decreases, with the LLC having the largest diameter and the pores the smallest (Fig. 2E-F, Fig. 3C, Supp. Mat. Table 3). A reduction in diameter was also observed when comparing the cross-sectional area of the LLC and primary tubules to the cumulative cross-sectional area within the same pore cluster, although primary tubules and pores clusters maintain similar values (Fig. 2E).

**Fig. 2.**
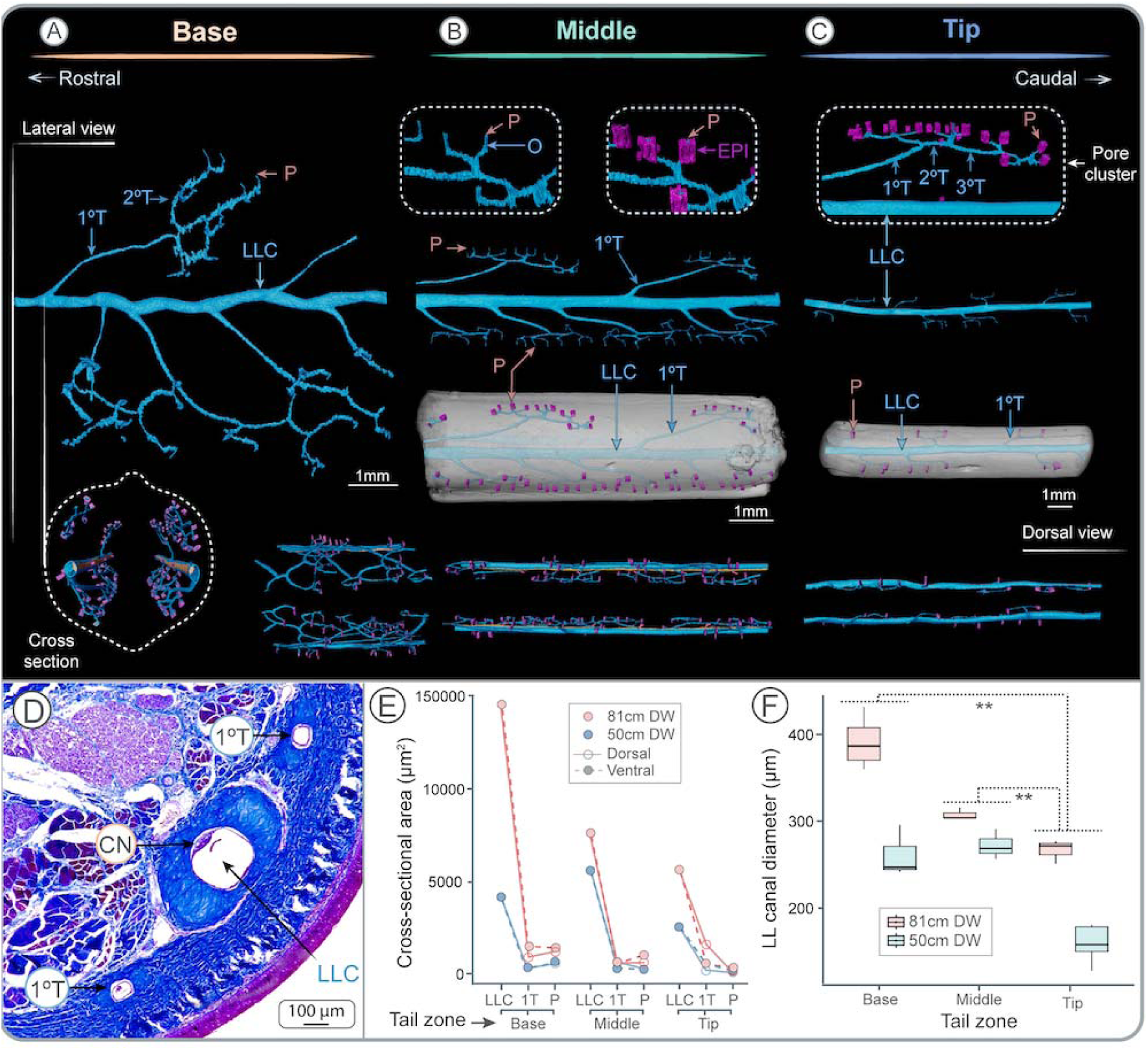
Lateral line morphology along the tail. **A–C)** µCT reconstructions of the lateral line canal, tubules and pores in the three zones of the tail showing the lateral line canal and tubules (blue) and pores (pink). The first tubules to divide from the lateral line canal are the primary tubules. Primary tubules continue dividing, forming secondary tubules which continue dividing until they reach the epidermis and open to the surrounding water via pores. Pores connected to a common primary tubule form a *pore cluster*. The cross-section (dashed lines represent the tail surface) and dorsal view show the position of both lateral line canals on each side of the tail. **D)** Mallory trichrome staining showing the lateral line canal, primary tubules, and the canal neuromast. **E)** Graphs showing variation in the cross-sectional area of the lateral line canal, tubules and the sum of the cross-sectional area of pores from the same pore cluster across tail zones, individuals and dorsal and ventral positions. **F)** Variation in lateral line canal diameter across tail zones and individuals. A double asterisk indicates that the differences were statistically different. **Abbreviations:** Branch **(B)**; Epithelium (EPI); Lateral line canal **(LLC)**; Pore **(P)**; Primary tubule **(1°T)**; Secondary tubule **(2°T)**; Tertiary tubule **(3°T)**; Opening **(O).**

**Fig. 3.**
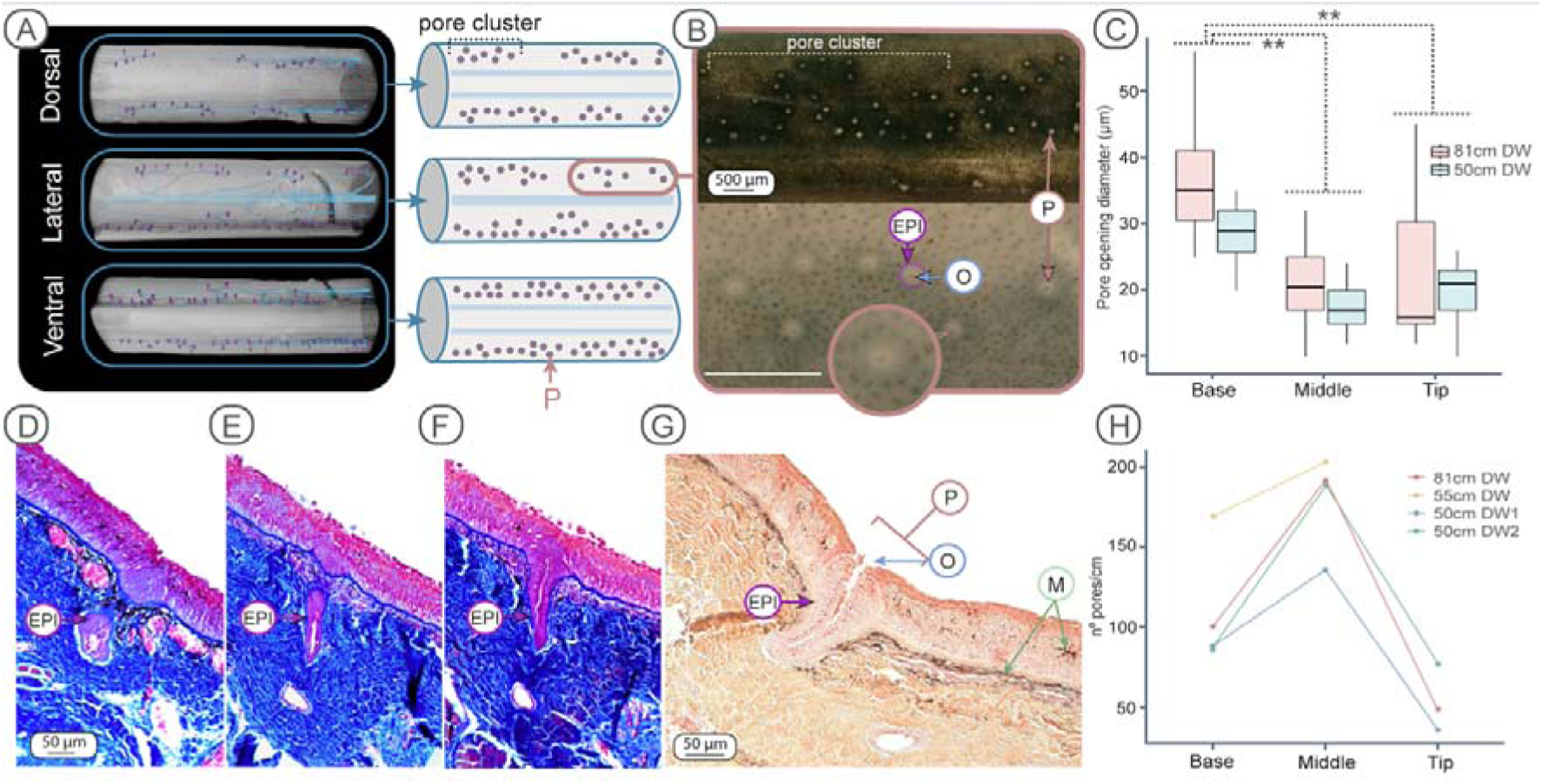
Pore distribution and morphology along the tail. **A)** µCT reconstructions and diagrams showing the pore distribution on the dorsal, lateral and ventral sides of the tail. Pore clusters consist of a group of pores that belong to the same primary tubule. Note the lack of pores on the lateral sides of the tail. **B)** Histological sections showing the pore distribution at the tail surface. Pores consist of an opening surrounded by an epithelial rim of columnar cells. **C)** Variations in pore opening diameters across tail zones for two individuals. Pores were significantly larger at the tail base, with no differences between the middle and the tip of the tail. Asterisks indicate significant differences. Pore opening diameters were proportional to the size of the animal. **D, E, F)** Mallory trichrome stained sections showing a pore opening in sequential sections. **G)** Hematoxylin-Eosin and Orange G histology sections showing the pore opening, where the lumen is lined by an epithelium. Note the melanocytes enclosed in the tail epidermis. **H)** Variation in pore density (n°pores/cm) across tail zones and among animals of different sizes. The tip for the 55cm DW individual was missing. **Abbreviations:** Epithelium **(EPI)**; Melanocytes **(M);** Pore **(P)**; Pore opening **(O).**

The lateral line system also varies in morphology, size and branching order across different tail zones. The diameter of the LLC decreases significantly from the base to the tip (*F* = 40.42, dF = 3.00, *P* < 0.01; Fig. 2F, Supp. Mat. Table 3) and it is positively correlated with the size of the animal, where larger animals have larger canal diameters (*F* = 40.42, dF = 3.00, *P* < 0.01; Fig. 2F). The number of tubule divisions from the LLC to the pores also varies across tail zones. The middle zone, which represents the larger proportion of the tail, had the greatest number of branches and the highest pore density (number of pores/tail cm) (Fig. 3H, Supp. Mat. Table 3). In contrast, the tip possesses the lowest number of divisions, particularly in smaller individuals, where the tubule network only presented two divisions until reaching the surface pores. This correlation and division pattern were similar for individuals of different sizes and sexes (Fig. 3H).

Pores are distinguished by the pore opening (area of the lumen of the tubule in contact with the surrounding water) enveloped by a thick rim of epithelial cells (Fig. 1A–B). Pores are distributed dorsally and ventrally following a linear pattern, but are not present at the lateral surfaces of the tail (Fig. 3A). Pore diameter (measured only considering the pore opening, Fig. 3B) varied across tail zones, having significantly larger diameters at the base, with no differences observed between the middle and the tip (Fig. 3C, F = 48.08, dF = 3.00, *P <* 0.01; Fig. 3C; Supp. Mat. Table 3). Similar to the LLC, pore diameter also increased with the size of the animal (*F =* 48.08, dF = 3.00, *P <* 0.01; Fig. 3C; Supp. Mat. Table 3). Both the LLC and tubules are surrounded by a thick layer of fibrous collagenous connective tissue, which becomes thinner as the canals and tubules decrease in diameter (Fig. 1F–H, Fig. 2D).

### (d) Neuromasts

Neuromasts were exclusively located in the LLC, and not observed in any of the tubules that lead to the pores. A combination of µCT scans, histology, and immunofluorescence showed that the canal neuromasts consist of a continuous structure along the entire tail, with not interruptions detected in any of our results (Fig. 4, Sup. Mat. Video 2). The continuity of the neuromast was particularly evident in PTA-stained and µCT-scanned samples, which allowed imaging of large tail sections (∼ 4cm). The PLLN run longitudinally along the tail beneath the canal (Fig. 4B). Nerve branches innervate the neuromast at regular intervals, always at constant distances between the division of two primary tubules (Fig. 4A–D, Sup. Mat. Video 2).

**Fig. 4.**
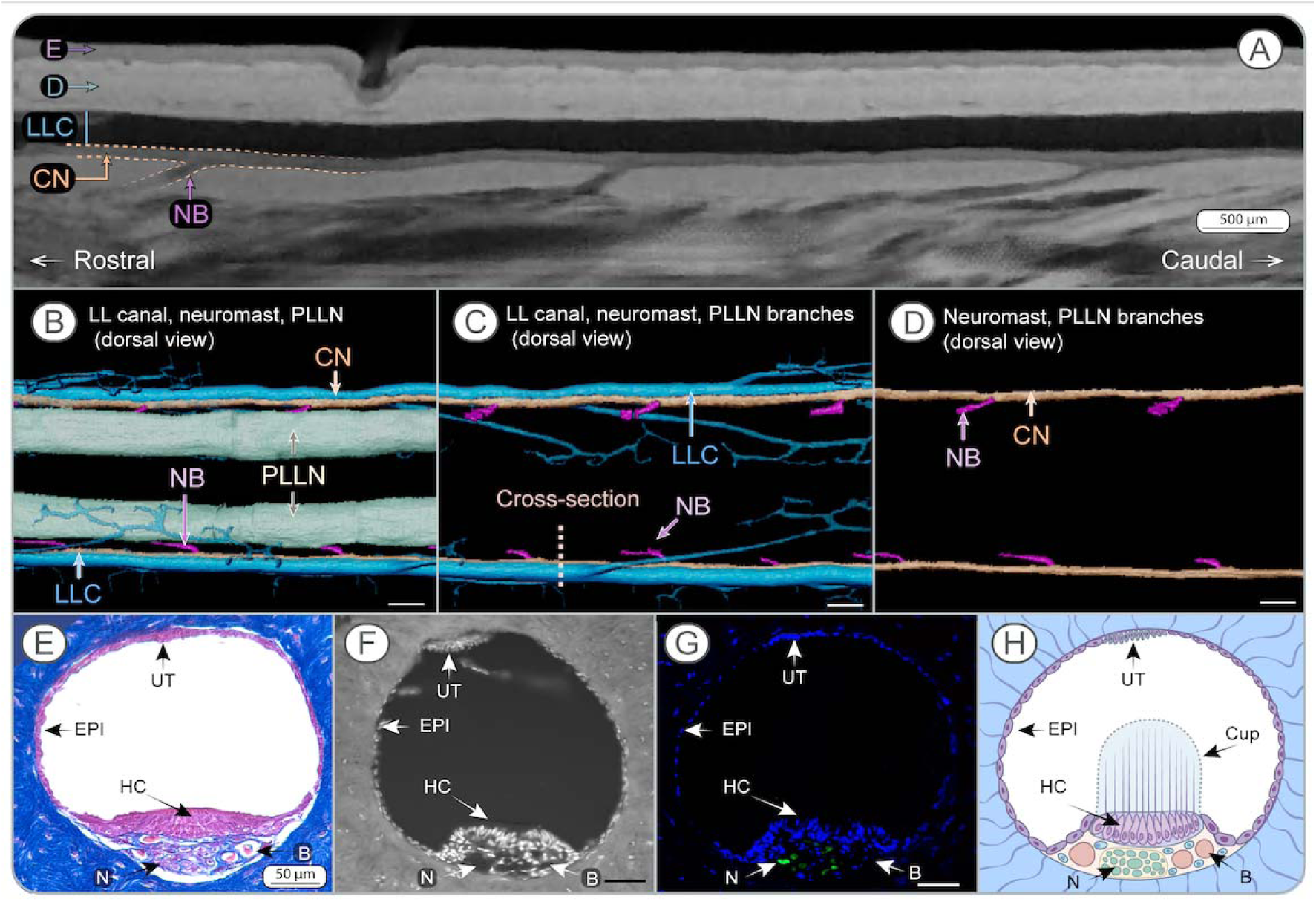
Anatomy of the continuous neuromast. **A)** µCT-scan of a tail stained with PTA showing the continuous neuromast enclosed in the lateral line canal, with nerve branches innerving the neuromast periodically. Dermis and epidermis are visible. **B–D)** 3D reconstruction from µCT scans showing the lateral line canals and tubules (blue), the continuous neuromast (orange), the nerve branches (pink), and posterior lateral line nerves that connect the neuromast with the spinal cord (green). Nerve branches are positioned between two primary tubules and connect the neuromast with the posterior lateral line nerves. Dashed line in C shows the cross-sectional orientation of images E–H. **F)** Mallory trichrome stained sections of the lateral line canal and the continuous neuromast. **E)** Confocal image with enhanced autofluorescence at 405nm showing the cellular nuclei (bright dots) and the surrounding dermis. **G)** Fluorescence image showing the cell nuclei DAPI stained (blue) and neural tissue stained with ß-III- tubulin antibody (green). **H)** Interpretation of the structural organization of the lateral line canal and the neuromast. Note that the cupula is based on an interpretation of the authors and was not able to be visualized in our sections. **Abbreviations:** Blood vessel **(B)**; Continuous neuromast **(CN)**; Cupula (Cup); Dermis **(D)**; Epidermis **(E)**; Epithelium **(EPI)**; Hair cells **(HC)**; Lateral line canal **(LLC)**; Nerve **(N)**; Nerve branch **(NB)** ; Nerve **(N)**; Primary tubule **(1°T)**; Posterior lateral line nerve **(PLLN)**; Unidentified tissue **(UT)**. Scale bars in E–G = 50 µm. Scale bars in B–D = 500 µm.

Histologically, the continuous neuromast is composed of sensory hair cells, identified by their central location in the neuromast structure (sensory zone), by their nucleus facing the basal surface of the cell, and by their organization in columns (Fig. 4F–H, Sup. Mat. Fig. 4). Blood vessels and nerves are visible running immediately beneath the hair cells along the LLC (Fig. 4F–H). An unidentified structure composed of a group of cells was found in the dorsal area of the LLC (opposite to the neuromast) (Fig. 4F–H). These cells stain with acid fuchsin (like epithelial cells) and did not test positive for β-III-tubulin, indicating the absence of nervous tissue in the structure. This structure was absent or appeared separated from the epithelium in most of the histology sections (e.g. Fig. 5A). Ciliary bundles of the hair cells and the cupula were not visible in any of our samples, mostly likely due to its loss during sample fixation.

**Fig. 5.**
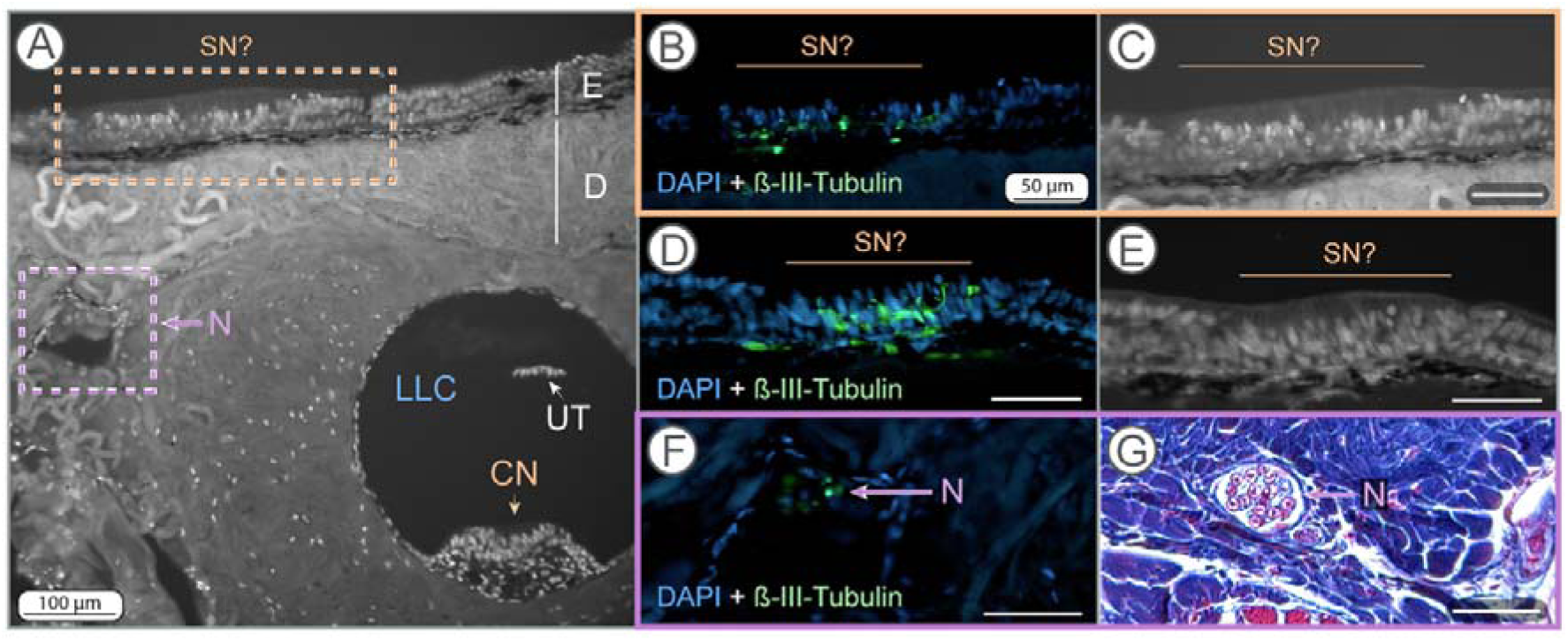
Putative superficial neuromasts. **A)** Confocal image obtained with the autofluorescence signal at 405nm showing the dermis with the lateral line canal containing the continuous neuromast, an ascending nerve, and the epidermis containing the sensory structures identified as potential superficial neuromasts. **B)** Sample stained with DAPI and ß-III-tubulin showing the cell nuclei and the nerve structures of the putative superficial neuromasts. **C)** Autofluorescence image of the putative superficial neuromast in B, showing the variation in cell morphology. **D)** DAPI and ß-III-tubulin staining of a different possible superficial neuromast. **E)** Autofluorescence image of D of similar variations in cell morphology. **F)** Nerve crossing the dermis stained with DAPI and ß-III-tubulin. **G)** Nerves crossing the dermis stained with Mallory trichrome. **Abbreviations:** Canal neuromast **(CN)**; Dermis **(D);** Epidermis **(E)**; Lateral line canal **(LLC)**; Nerve **(N)**; Potential superficial neuromast **(SN?)**; Unidentified tissue (UT). Scale bars in B–G= 50 µm.

Immunofluorescence showed sensory structures embedded in the epidermis positive for β-III-tubulin (Fig. 5). These structures, which were uncommon and exclusively found in the epidermis, are located parallel to the LLC and cover areas devoid of pores (Fig. 5A). These structures were not located in pits and presented a different cell morphology to the surrounding epidermal cells and we identify them as putative superficial neuromasts (Fig. 5B–E). Exclusively in this region, nerves bordered the collagenous dermis surrounding the LLC and ran toward the epidermis (Fig. 5F–G).

## 4. Discussion

Using a combination of different imaging techniques, this study describes the complex anatomical systems within the tail of *Rhinoptera bonasus*, the cownose ray. Our results showed that the tail is a long and slender structure with a complex internal anatomy. The tail is characterized by having a vertebral column with an unusual morphology and by the presence, at both sides of the tail, of a lateral line system. The lateral line system is formed by a lateral line canal (LLC) that contains a continuous neuromast and is connected to the environment through a highly branched network of tubules. Analyses of tail morphology in other myliobatid species confirms that these features are not unique to cownose rays. To facilitate understanding the results of this study, in Figure 6 we present a graphical summary of the main morphological results and further interpretation and hypothesis of the lateral line stimulus filtering.

**Fig 6.**
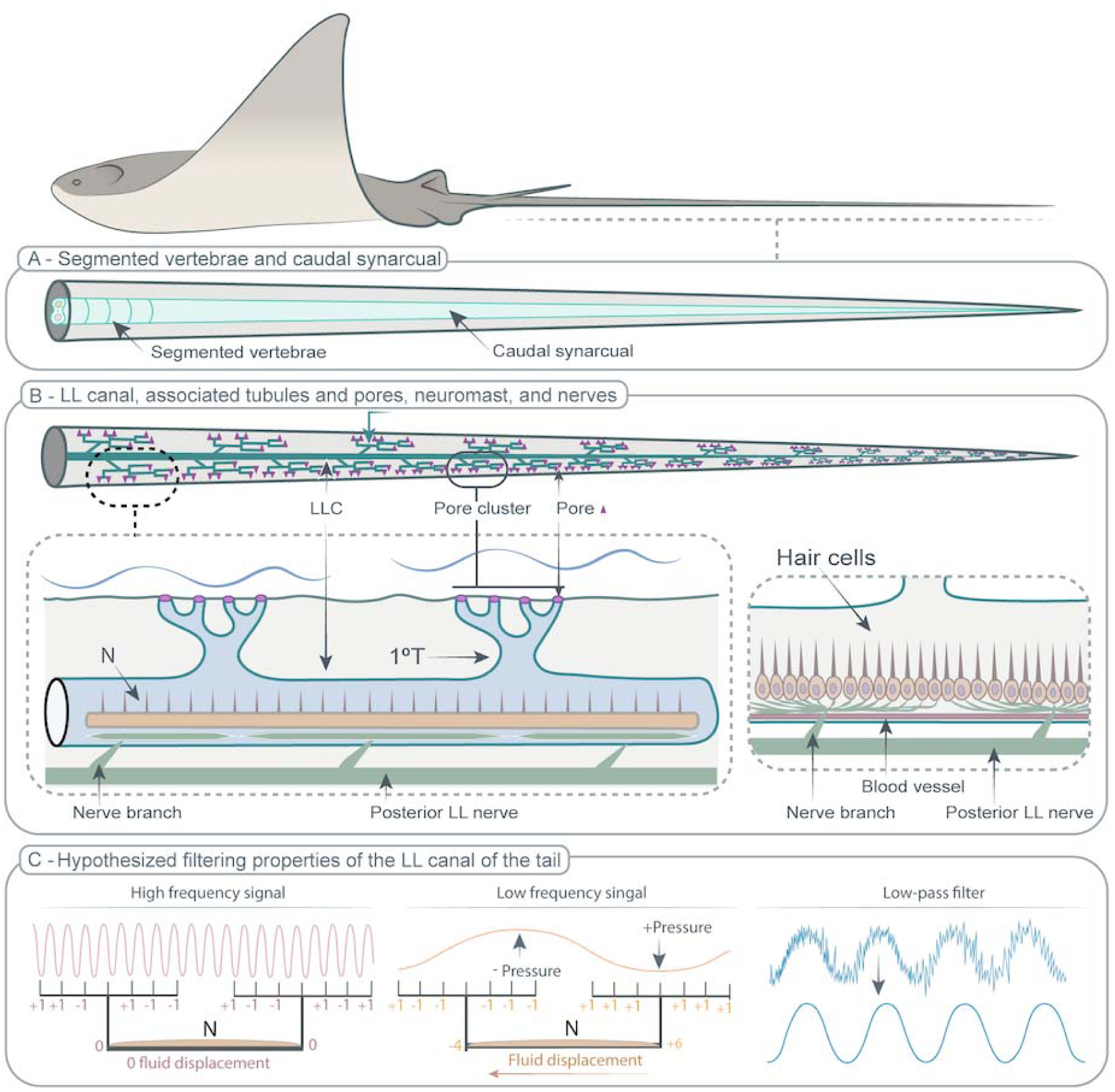
Anatomy and hypothesized function of the tail of *Rhinoptera bonasus*. **A)** Vertebral morphology variation along the tail. At the base, the vertebral column is formed by separated and well-defined vertebral units that allow active control of the tail. Caudally, vertebrae are fused and form the caudal synarcual which provides passive stiffness to the tail. **B)** The lateral line canal, with branched tubules that open to the surrounding water via pores. The lateral line canal contains the continuous neuromast. The continuous neuromast is connected via nerve branches to the posterior lateral line nerve. **C)** The hypothesized filtering properties of the lateral line canal of the tail. The spatial distribution of pores would allow filtering high- frequency signals (low-pass filter), while the increase in diameter from the pores to the lateral line canal would allow filtering low-pressure signals (and, therefore, low amplitude).

### (a) Tail gross anatomy

As in other vertebrates, the tail of *R. bonasus* is an extension of the vertebral column but, instead of being formed by sequentially interconnected vertebrae, the vertebral column presents significant variation along the length of the tail. At the base (between the pelvic fins and the posterior end of the barb) vertebral units are organized like a closing zipper (like other parts of the body, [21]) and are attached to a substantial musculature. However, from the posterior end of the barb to the tip, the muscular content is reduced, and the vertebral column is reshaped into a fused mineralized rod forming a continuous mineralized structure not divided by vertebral segments (Fig. 6A). This fused structure is similar to the synarcual described in other batoid [22, 23].

In chondrichthyans (especially chimaeras and batoids) the synarcual is defined as that portion of the vertebral column with fused segments (similar to the sacrum and pygostyle in tetrapods) [22–25]. In batoids, the synarcual within the body is a synapomorphy (i.e. shared derived feature of this clade) [26]. It is located in the anterior part of the vertebral column connected to the chondrocranium and expanding beyond the pectoral fins [22, 27]. Due to its plate-like morphology, the synarcual has a biomechanical function, as it gives support to the enlarged pectoral fins [3, 23, 28]. Myliobatiforms are the only batoid group that has a second thoracolumbar synarcual located in the posterior area of the body (posterior to the pectoral fins), although its function is not known [26, 29]. The fused caudal vertebrae structure of cownose rays’ tail is similar in several ways to the synarcual described in other batoids, including being a continuous fused skeletal structure covered by tesserae, lacking vertebral centra, and containing a space for the spinal cord and the hemal canals that transverse the mineralized tissue through foramina. Therefore, we propose the term ‘caudal synarcual’ for the fused vertebrae structure of the *R. bonasus* tail.

Variation in skeletal morphology along the tail suggests variation in tail flexibility and mobility. The articulated vertebrae in the base zone of the tail, in combination with the larger mass of tail musculature located here, likely allow overall tail mobility, support (the tail is most often held horizontally behind swimming rays, and does not sag ventrally), and orientation of the entire tail structure. In contrast, the caudal synarcual increases passive stiffness in the middle and distal tail regions and, in combination with the reduced muscle mass, should limit mobility and allow little deformation of the distal half of the tail. Therefore, our anatomical data suggest that cownose rays could control overall tail orientation from the base, while the more distal stiff tail region would be relatively incapable of independent bending or undulation (Fig. 6A). This result suggests that the tail is not an active propulsive structure contributing to thrust generation during locomotion, which contrasts with other slender tails, such as the tapered tails of lizards or the caudal region of fishes which are articulated and mobile along their length [12, 30]. The mineralized structure of the *R. bonasus* tail may be important for increasing tail stiffness that will enable the tail to remain elevated to avoid contact with the substrate during foraging, to assist in maintaining stability while swimming, and importantly as a stable platform for the remarkable lateral line sensory system located within the tail.

### (b) The tail sensory system

In teleost fishes, the lateral line system (comprising the LLC, tubules, and neuromasts) is mechanosensory, as it detects water movements that cause pressure gradients among pores, moving the fluid that fills the tubules and canals [31, 32]. This fluid movement causes a displacement of the cupula that covers neuromasts, producing a deflection of the kinocilia and stereocilia of the hair cells and, ultimately, informing the central nervous system of the stimuli [31, 33, 34]. The presence of the LLC with neuromasts indicates that the tail of *R. bonasus* is a mechanosensory structure, able to detect hydrodynamic stimuli generated by movements of the surrounding water (such as movements generated by nearby rays, a predator, prey items, or the body of the ray itself) (Fig. 6B).

Structurally, the lateral line system of the tail of *R. bonasus* contrasts with the posterior lateral line system (lateral line system found in the trunk and tail regions) described in sharks and teleost fishes [9, 35, 36]. In sharks and teleosts, the posterior lateral line system is characterized by a LLC that opens to the lateral sides of trunk and tail through short, unbranched tubules [36]. This differs from the highly branched lateral line of *R. bonasus* tail, which opens to the dorsal and ventral surface of the tail (Fig. 6B) [37]. However, highly branched lateral line systems have been described in the body of other stingrays [6, 7, 9], which may be an indicative of the importance of a branched lateral line systems in the biology of batoids from the order Myliobatiformes. In *R. bonasus*, the lateral line tubules and canals are surrounded by a collagenous layer that varies in thickness proportional to canal diameter. Similar to the tunica media in circulatory system [38], this thickness variation may indicate a mechanical role in maintaining the shape of the canals and tubules. Our results also show a group of cells located opposite to the neuromast in the LLC, which have not been previously described in other lateral line systems. It was not possible to determine whether these cells form a continuous or discrete structure within the LLC and, while the precise function of these cells remains unclear, our results indicate that they do not contain nervous tissue.

In the natural environment, hydrodynamic stimuli produced by biotic and abiotic sources are highly variable and include surface waves, local currents, prey, predators, conspecifics and body deformations [39, 40]. Some of these stimuli could be considered environmental noise (unwanted water movements that interferes with a relevant specific stimulus) and the lateral line structure can filter these unwanted stimuli enhancing detectability of specific frequencies and amplitudes [31, 41, 42]. The filtering properties are determined by the structure of the neuromast cupula as well as the morphology of the LLC and tubules [43, 44]. We suggest that differences in branching pattern, cross-sectional area, pore distribution and pore density determine the sensitivity and filter properties of the lateral line system in the tail of *R. bonasus* (Fig. 6C) [7, 41–44].

Based on Laplace’s law and simulations correlating lateral line morphology and filter properties [41, 42], the reduction of cross-sectional area from the LLC to the pores indicates that high pressures are required at the pore surface to move the lateral line fluid within the LLC (resulting in a deflection of the cupula and, thus, the hair cells of the neuromasts). As the diameter of the tail lateral line system is reduced from the LLC to tubules to the pores, the effective Reynolds number will be reduced which reflects the corresponding increase in viscosity of the fluid and also act to filter external signals. Since the pressure of a wave is correlated with its amplitude, this suggests that the tail lateral line system may function as a high-amplitude pressure detector. The lack of non-pored tubules (pores not connected to the surface) suggests that the *R. bonasus* tail does not function as a tactile sensor. This result contrasts with the presence of non-pored tubules described in the body of other batoids and the rostral saw of sawfishes, where non-pored tubules have been interpreted to give a tactile function to detect buried prey [7, 45].

The sensitivity of the lateral line system is determined by a combination of factors, including the tree topology of the network and the density and spatial distribution of pores [7, 41, 42]. A highly branched network of tubules allows an increased number of pores to connect the neuromasts to the surrounding water. An increased number of pores expands the area capable of detecting hydrodynamic stimulus, allowing *R. bonasus* to perceive water movements along the entire length of its tail [41].

The distribution of the surface pores can determine which frequencies are being filtered [41, 42]. Each pore cluster on the tail of *R. bonasus* contains around 15-20 pores arranged linearly and closely spaced. The pressure differences among pores within the same cluster should be averaged when reaching each primary tubule. When the wavelength of a given stimulus is similar to the distance between two pores, the pressure difference in the tubule bifurcation will be zero, resulting in no fluid displacement (Fig. 6C). Conversely, stimuli with lower frequencies will produce a pressure gradient between two different primary tubules, causing a fluid displacement within the canal where the neuromast is located. The averaging of pressure differences in the tubules could also allow them to function as spatial filters, attenuating the transmission of high-frequency disturbances and emphasizing lower frequency stimulus. Thus, the lateral line system in the tail of *R. bonasus* could function as a low-pass filter (Fig. 6C).

No differences were observed in the pattern and distribution of pores between sexes. However, variations related to lateral line diameter, animal size and tail zone were observed. First, the lateral line system (including canal, tubules and pores) appears to scale with the size of the animal, although ontogenetic studies are needed to verify this result. For each tail zone along the length of the canal (base, middle, tip), the middle region of the tail contains the highest number of branching tubules and, consequently, the highest density of pores. This suggests that this area is more sensitive than the tail base and tip.

### (c) The continuous tail neuromasts

We have demonstrated that the neuromasts within the LLC present a *continuous* structure along the entire length of the tail (Fig. 6B). This continuous neuromast appears to be composed of a continuous epithelium with numerous hair cells, as no breaks in the epithelium were observed during the analysis of any of our samples. However, due to the difficulty of imaging the gelatinous cupula (if present), it is not known whether the cupula also represents a continuous structure. This continuous neuromast differs from what is known in teleost fishes, where neuromasts are discrete structures located between the division of primary tubules that extend to the skin surface [34, 37]. Long neuromasts have been diagrammed once previously in elasmobranchs, although direct demonstration of the continuity of neuromasts has not been previously shown, especially over such an extensive distance as described in the tail of *R. bonasus* [2, 6, 46]. In contrast, discrete neuromasts have been reported in LLC of the head in embryos of the little skate (*Leucoraja erinacea*) [47]. The difference in neuromast length (continuous vs. discrete) between *R. bonasus* and *L. erinacea* may be related to three possible options: 1) neuromasts may grow in length with ontogeny, forming a continuous structure in adult batoids; 2) neuromast length may vary across different body regions in both rays and skates; or 3) neuromast length may differ among batoid species. Further research is needed to explore these hypotheses.

The cross-sectional anatomy of the neuromasts in *R. bonasus* shows that they contain hair cells (sensory units) in a similar position and morphology to neuromasts in teleost fishes. Posterior lateral line nerves innervate the neuromast through nerve branches at period distances between the division of two primary tubules. This confirms the hypothesis proposed by [46] where, although being a continuous structure, hair cells in a continuous neuromast are divided into packages or groups, with the information collected by these cell groups is transported by a specific nerve branch (Fig. 6B). This division allows the information to be localized spatially, so the animal may be able to recognize where, along the LLC, an hydrodynamic disturbance may be located. Further studies are needed to understand the function and advantages given by continuous neuromasts and if this is a conserved structure across other species of chondrichthyans.

We also observed putative sensory structures in the tail epidermis. These structures were uncommon and only visualized using immunofluorescence. Although the cupula was not preserved, we believe that these structures are superficial neuromasts. Superficial neuromasts have been described in the tails of myliobatiforms such as *Dasyatis sabina* [6] and *Myliobatis australis* [48] enclosed in pits and termed pit organs. However, the putative superficial neuromasts found in this study do not occur in pits, but are structures embedded in the epidermis. These structures are also not associated with the small tail denticles (identified as claw type prickles, [49]), whose function remains unknown. These superficial neuromasts in the cownose ray tail were scarce and distributed on the skin parallel to the LLC, in the area that lacks lateral line pores. This agrees with the distribution of superficial neuromasts previously described in other batoid tails [6, 50, 51].

## 5. Conclusion

Myliobatid stingrays are characterized by having a long and slender tail that extends behind the body. A tail with this morphology is characteristic of the group and is not used to generate thrust during locomotion. We used a combination of imaging techniques to describe the anatomy of the tail in *Rhinoptera bonasus* (the cownose ray) and our results show that the tail is stiffened due to the presence of a mineralized rod. This likely increases tail stability by passively damping perturbations during swimming as well as reducing hydrodynamic noise by limiting tail movement. The tail also contains a lateral line system with a complex structure, formed by two lateral line canals on each side of the tail which contain continuous neuromasts connected to an extensive tubule network. This branched tubule network is proposed to act as a filter, ultimately influencing the hydrodynamic stimulus received by the neuromasts. Therefore, we hypothesize that this network filters low-amplitude, high-frequency hydrodynamic stimuli. Cownose rays are active swimmers living in a diversity of environments where background noise can be high: for example, noise produced by waves near the surface or during swimming (signals associated with high frequencies). The filtering capacities of the lateral line system of the tail could improve the signal-to-noise ratio, avoiding overstimulation of the canal neuromasts. We propose that the tail of *R. bonasus* may act as a hydrodynamic antenna able to detect water disturbances resulting from prey, predators, body movements, and near body flow dynamics.

## Supporting information

Sup. Mat. Figure 1

Sup. Mat. Figure 2

Sup. Mat. Figure 3

Sup. Mat. Figure 4

## Ethics

This work did not require ethical approval from a human subject or animal welfare committee.

## Data accessibility

Data is available from the authors upon request.

## Declaration of AI use

No AI-assisted technologies were used in creating this article.

## Author’s contributions

J.C.: conceptualization, funding acquisition, data curation, formal analysis, anatomical analyses, visualization, writing—original draft, writing—review and editing; G.V.L.: writing—review and editing; funding acquisition, conceptualization, project administration, resources. Both authors gave final approval for publication and agreed to be held accountable for the work performed herein.

## Acknowledgments.

Authors thank Connor F. White for the insightful discussion on results interpretation and assistance with R coding, particularly using ggplot. We also thank Jacqueline Webb, Matt McHenry, and Huntang Ko for their contributions to the interpretation of results, to Tim Handsel and Benjy Davis (Ripley’s Aquarium of Myrtle Beach) and Tonya Wiley (Havenworth Coastal Conservation) for the samples, to Marina Rodriguez and Nicolás Romero for their advice and support in the application of the immunofluorescence protocols, and to Gayani Senevirathne for her assistance with the confocal microscope.

## Supplementary Material

Due to size limitations in the BioRxiv submission interface, videos in the supplementary materials can be downloaded from the following Dryad link: http://datadryad.org/stash/share/-5EVWSD7Qjbo6ogRIuNQhZwpvKA8h38qYGe_5CzGFq4

**Figure 1.**
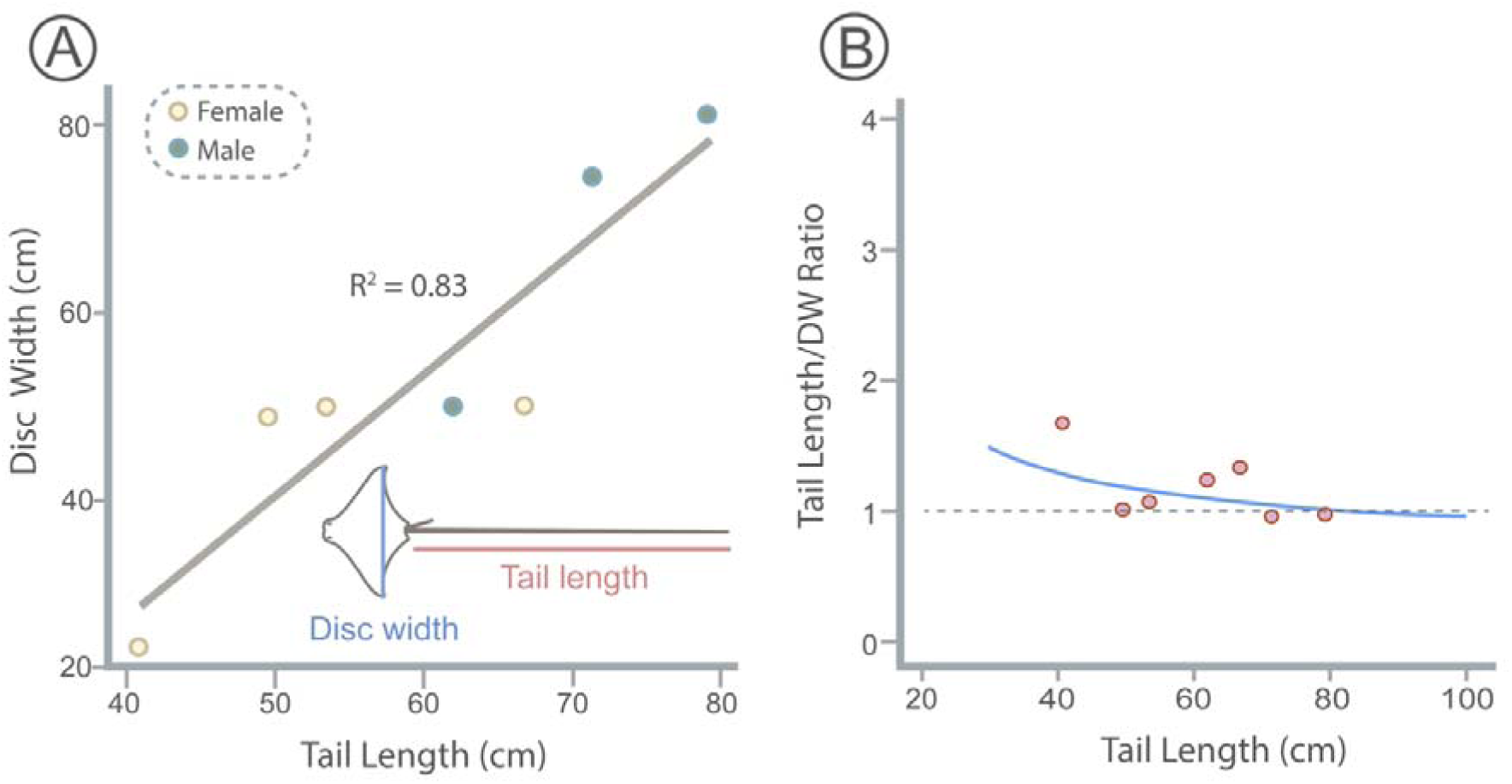
Scaling of tail and body size in the cownose ray, *Rhinoptera bonasus*. **A)** A linear regression showing, for all individuals measured (including males and females, Sup. Mat. Table 1), how the tail length significantly increases with the disc width (DW) (*P* = 0.004, *R^2^* = 0.83). **B)** Tail length / DW ratio is negatively correlated with tail length and based on the linear regression in A, this relationship is non-linear (blue curve). This implies that smaller individuals have proportionally longer tails, which is supported by the fact that the smallest individual used in this study has a tail 1.7 times larger than its DW.

**Figure 2.**
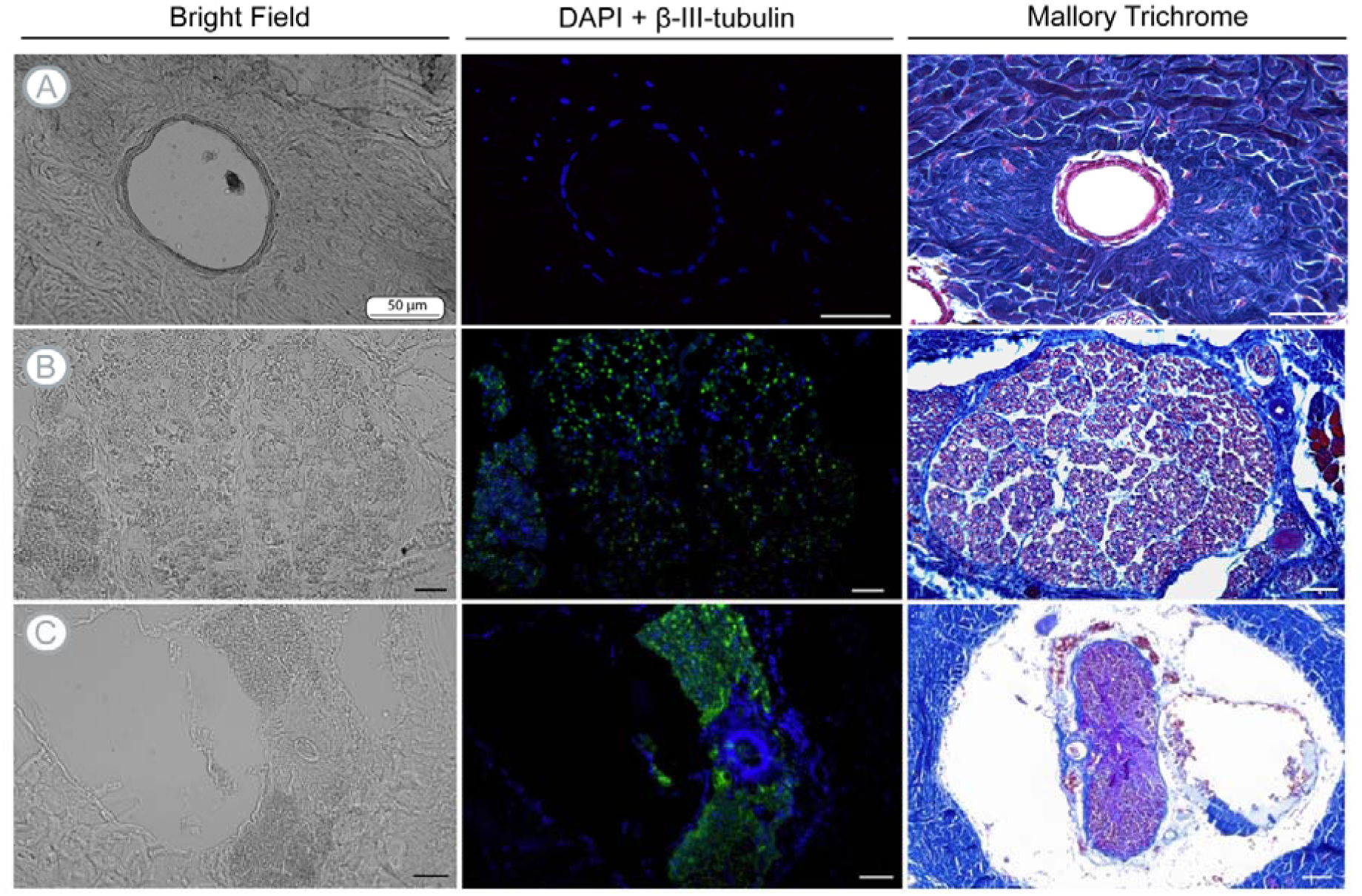
Immunofluorescence controls imaged using bright field, DAPI and ß-III-tubulin, and Mallory trichrome. **A)** Tissue composition of the lateral line tubule, showing the epithelial tissue surrounding the lumen and the lack of nervous tissue. **B)** A cross-section of the posterior lateral line nerve, positive for ß-III-tubulin (green). **C)** Cross-section of the spinal cord, positive for ß-III-tubulin (green). Posterior lateral line nerves and spinal cord were used as ß-III-tubulin antibody controls.

**Figure 3.**
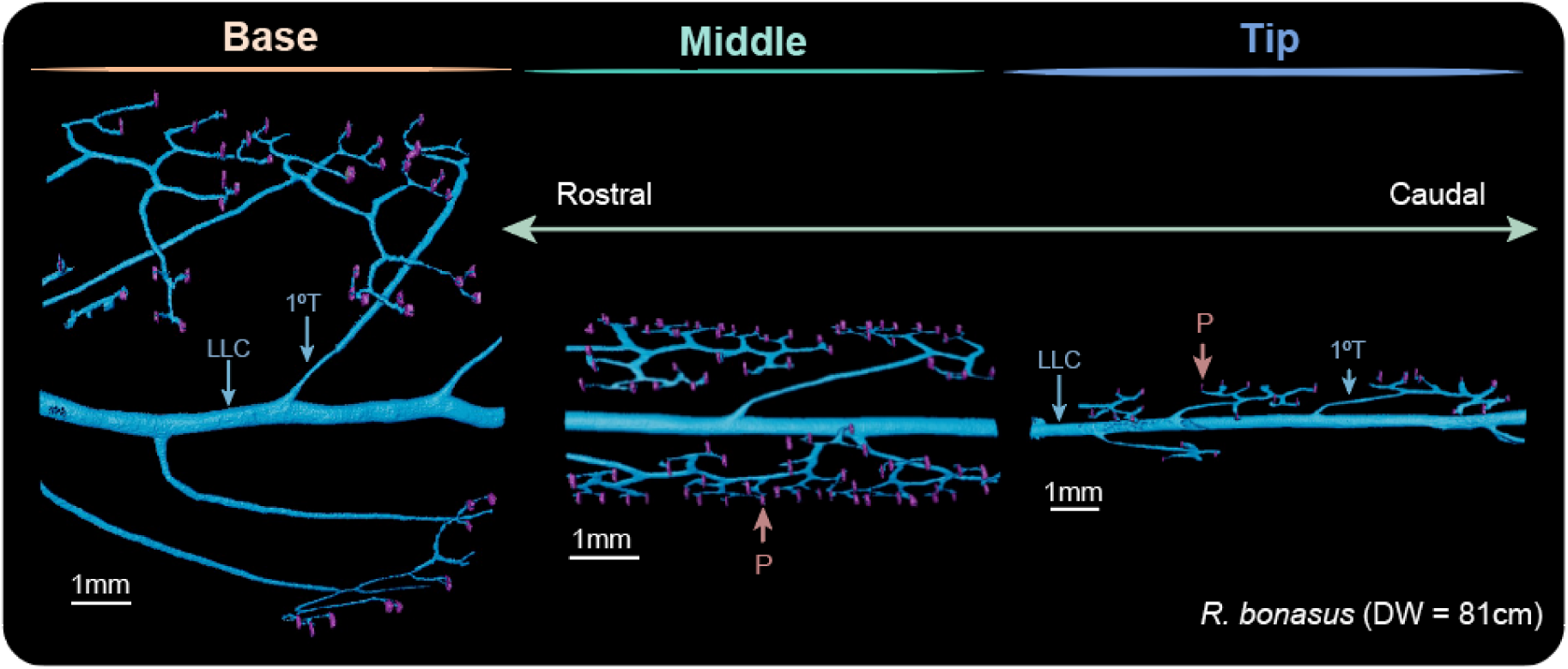
Lateral line morphology of *Rhinoptera bonasus* (DW = 81cm). Lateral line morphology variations across different tail zones (base, middle and tip). Note the reduction in lateral line canal diameter from the base and the tip (caudally), as well as the higher number of divisions and pores of the middle zone of the tail. **Abbreviations:** Lateral line canal (LLC); Primary tubule (1°T); Pore (P).

**Figure 4.**
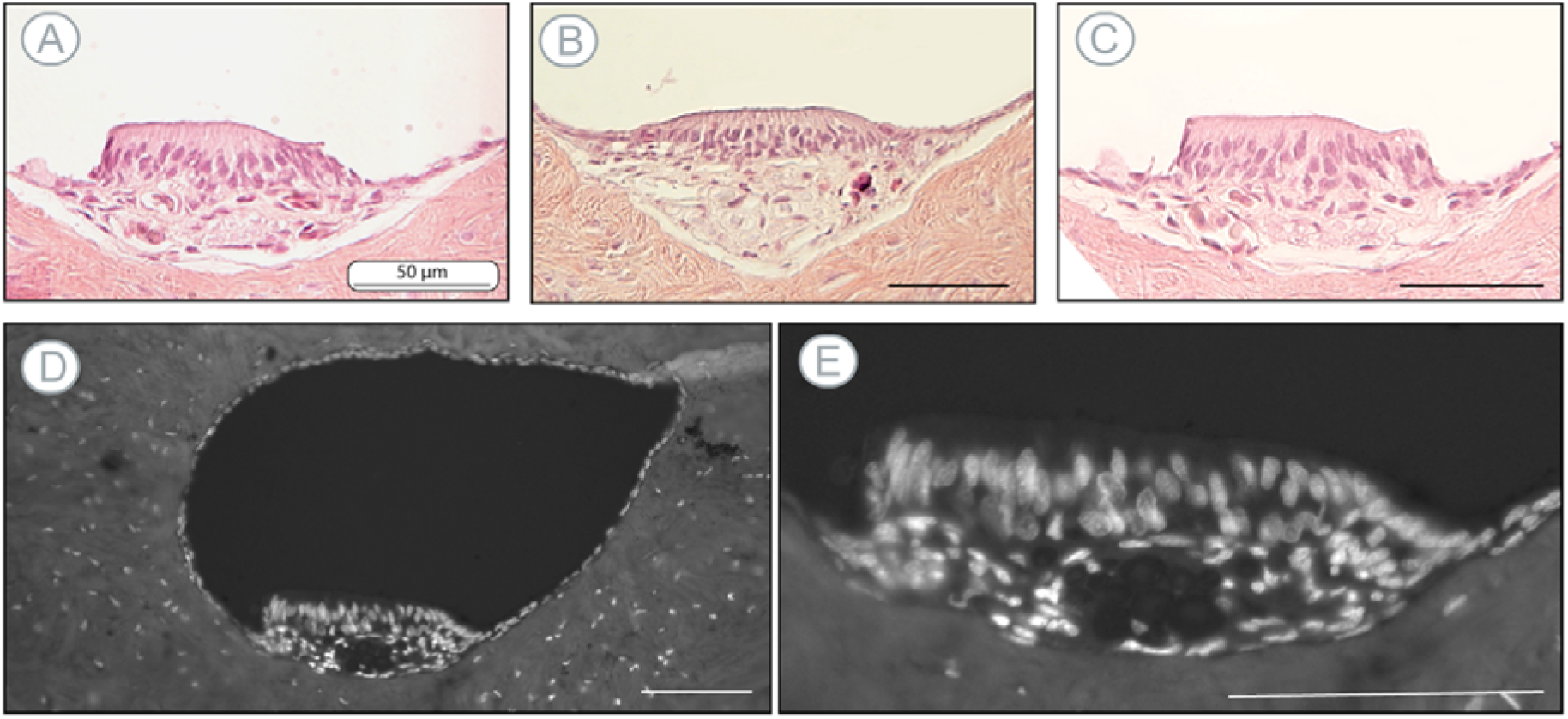
Cross-section of the continuous neuromast. **A–C)** Neuromast cross-sections stained with hematoxylin-Eosin and Orange G showing the position of the cellular nucleus. Note the basal orientation of the hair cell nucleus, located at the base of the neuromast. **D–E)** Confocal images obtained using DAPI staining and autofluorescence excited at 405nm. Note the difference in the position and size of the nucleus of different cells.

**Table 1.** *Rhinoptera bonasus* specimens used in this study. with the morphometrics measured (DW = disc width), sex, and analysis performed for each individual. The specimens with the MCZ belong to the specimens stored in 70% ethanol at the ichthyology collection of the Museum of Comparative Zoology of Harvard University.

| Specimen | DW (cm) | Tail length (cm) | Sex | Analysis |
| --- | --- | --- | --- | --- |
| R. bonasus_florida | 50 | 62 | Male | Morphometrics, pore analysis, μCT-scan, PTA staining, histology, immunofluorescence |
| R. bonasus_aquarium | 81 | 79 | Male | Morphometrics, μCT-scan, PTA staining, histology, immunofluorescence |
| R. bonasus_MCZ_49097 | 50 | 53 | Female | Morphometrics, pore analysis, μCT-scan |
| R. bonasus_MCZ_535_2 | 50 | 66 | Female | Morphometrics, pore analysis |
| R. bonasus_MCZ_317 | 24 | 40 | Female | Morphometrics, pore analysis |
| R. bonasus_MCZ_374 | 74 | 71 | Male | Morphometrics, pore analysis |
| R. bonasus_MCZ_535_1 | 48 | 50 | Female | Morphometrics, pore analysis |
| R. bonasus_MCZ_746 | 55 | Tail tip missing | Male | Pore analysis, pore analysis |

**Table 2.** Measured variables for different samples. Abb: Lateral line canal (LLC)

| Specimens and structures | Variables |
| --- | --- |
| Specimen | Disc width (cm), body length (cm), total length (cm) |
| Tail | Length (cm), diameter (cm) |
| Lateral line | Diameter of each branch level ($\mu\text{m}$), cross-sectional area ($\text{cm}^2$)<br>distance between 1<sup>st</sup> level branching, number of branches, branching<br>level division, angle among LLC and primary tubules, ratio among<br>LLC and primary tubules diameter (diameter LLC canal / diameter<br>primary tubule) |
| Pores | Pore diameter ($\mu\text{m}$), cross-sectional area ($\text{cm}^2$), pore density (n°pores<br>/ 1cm length) |

**Table 3.** Lateral line measures. Abb: Lateral line canal (LLC)

| Specimen | Tail zone | Tail position | LLC diameter (µm) | x □ Pore diameter (µm) | Σ Pore diameter (µm) – per cluster | N° Pores per cluster | Σ cross-sectional area LLC (mm²) | Σ cross-sectional area pores (mm²) |
| --- | --- | --- | --- | --- | --- | --- | --- | --- |
| *R. bonasus*<br>(DW = 81cm) | Base | Dorsal | 431 | 33.23<br>±6.3 | 433 | 13 | 0,14 | 0,011 |
|  |  | Ventral | 431 | 41.4<br>±7.7 | 414 | 10 | 0,14 | 0,013 |
|  | Middle | Dorsal | 311 | 19.7<br>±4.7 | 350 | 17 | 0,07 | 0,005 |
|  |  | Ventral | 311 | 23.72<br>±6.8 | 495 | 22 | 0,07 | 0,005 |
|  | Tip | Dorsal | 179 | 31.75<br>±1.7 | 36 | 5-6 | 0,02 | 0,0005 |
|  |  | Ventral | 179 | 18.00<br>±8.6 | 92 | 4 | 0,02 | 0,0016 |
| *R. bonasus*<br>(DW = 50cm) | Base | Dorsal | 229 | 29.12<br>±3.2 | 233 | 8 | 0,04 | 0,005 |
|  |  | Ventral | 229 | 28.33<br>±4.7 | 273 | 4-6 | 0,04 | 0,005 |
|  | Middle | Dorsal | 268 | 17.60<br>±2.00 | 176 | 7-8 | 0,05 | 0,002 |
|  |  | Ventral | 268 | 17.93<br>±3.63 | 142 | 10 | 0,05 | 0,002 |
|  | Tip | Dorsal | 267 | 17.33<br>±3.9 | 94 | 2-4 | 0,05 | 0,0011 |
|  |  | Ventral | 267 | 23.12<br>±2.23 | 127 | 2 | 0,05 | 0,0031 |

**Video 1. 3D reconstruction of the lateral line of *Rhinoptera bonasus* (DW=50cm).**

**Video 2. µCT scan of the lateral line of *Rhinoptera bonasus* (DW=50cm) showing the continuity of the neuromast.**

