## Supplementary figures and images for "A hydrodynamic antenna: novel lateral line system in the tail of myliobatid stingrays"

### Sup. Mat. Figure 1

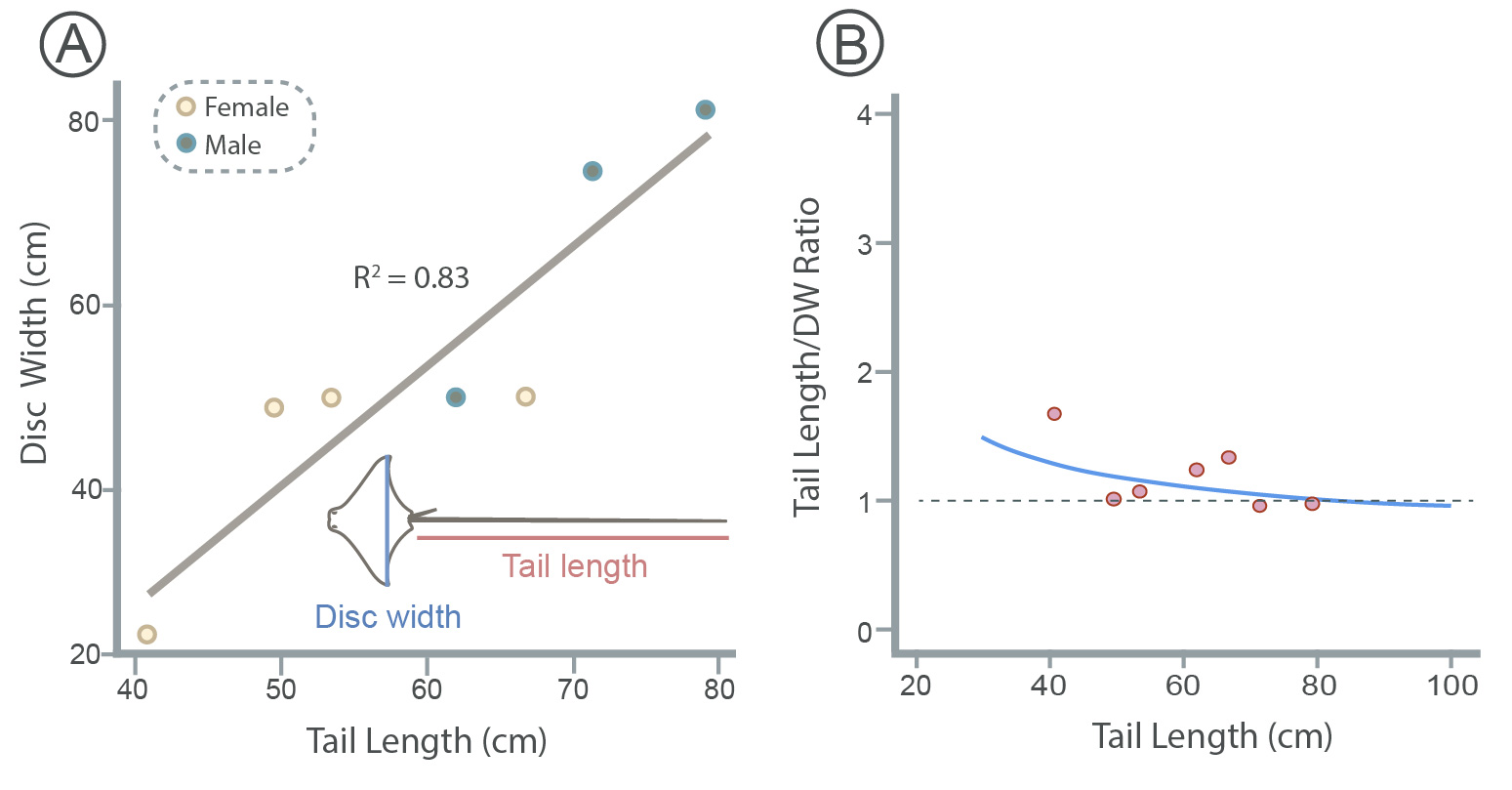

### Sup. Mat. Figure 2

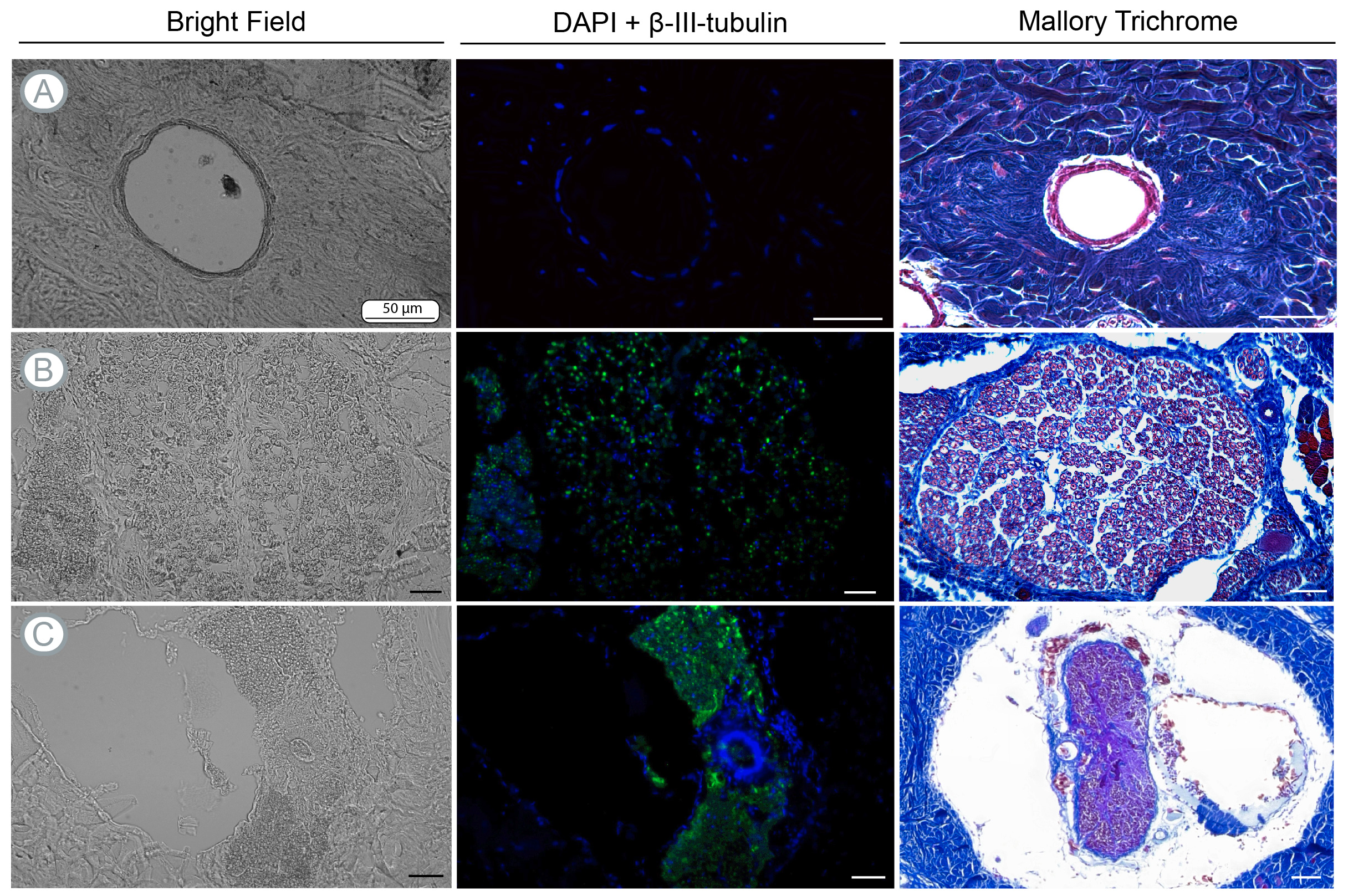

### Sup. Mat. Figure 3

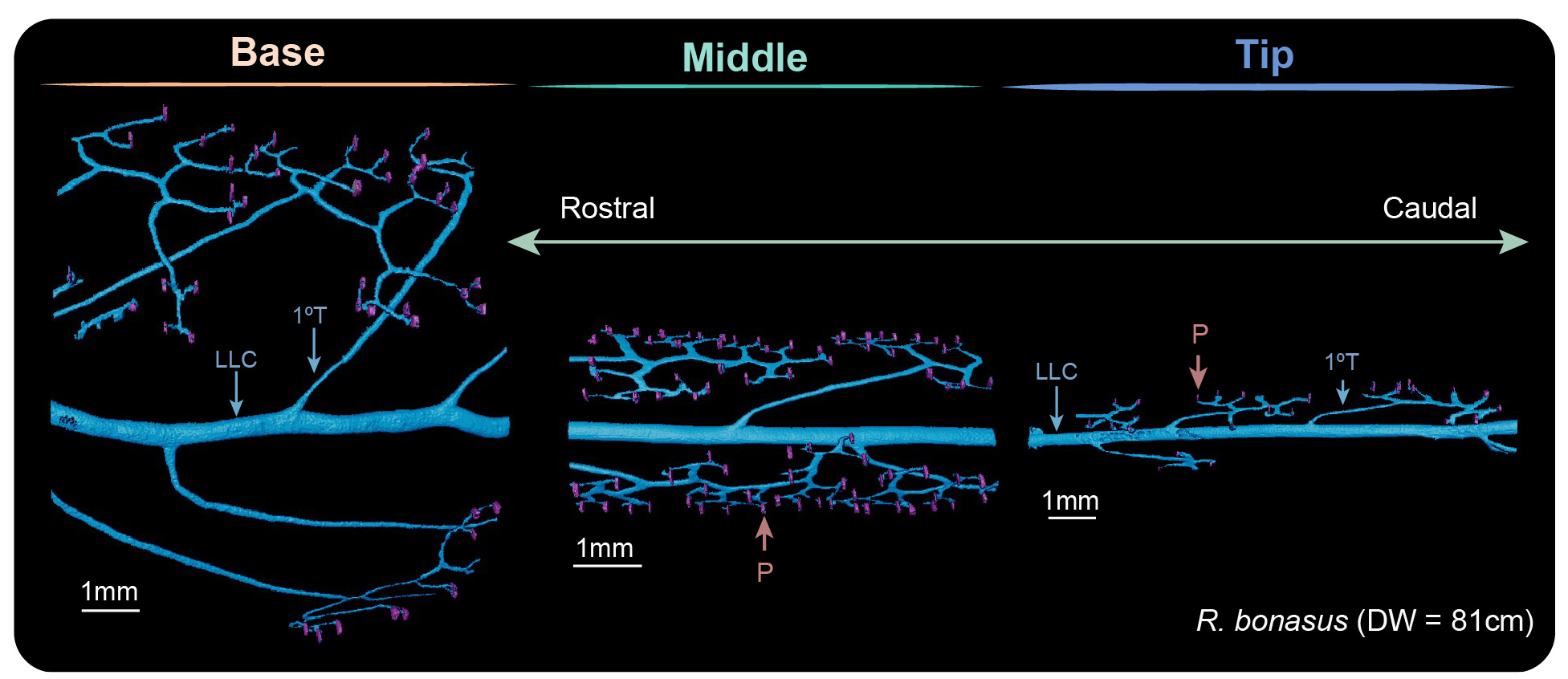

### Sup. Mat. Figure 4

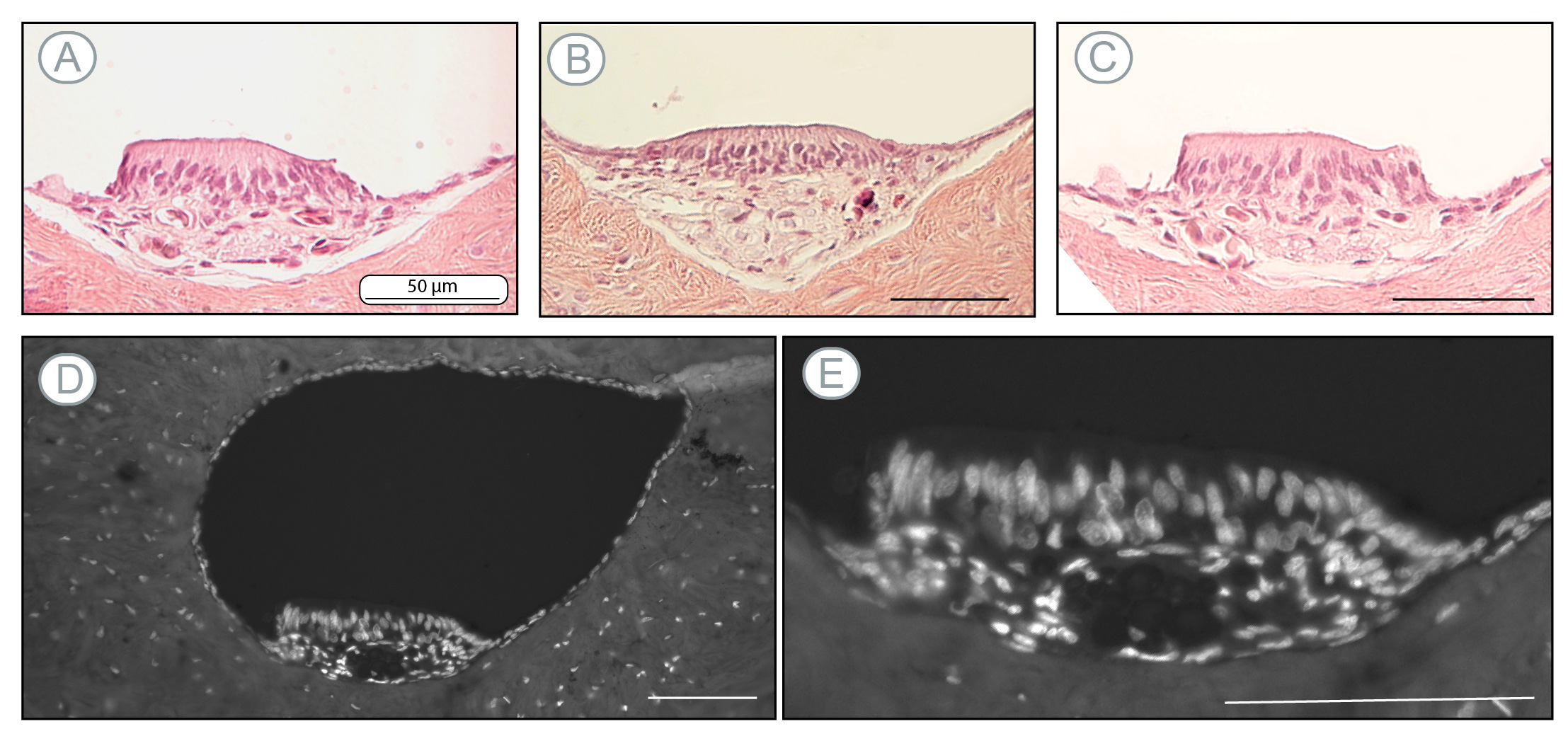
